# Let-them-stick: Increasing biofilm formation by the acetogen *Sporomusa ovata* through adaptive laboratory evolution

**DOI:** 10.1101/2025.06.26.661873

**Authors:** Louise V. Grøn, Laura Munoz-Duarte, Ian P.G. Marshall, Niels Krabbe Johnsen, Bekir Engin Eser, Michael W. V. Kofoed, Klaus Koren, Jo Philips

## Abstract

Acetogenic bacteria are attractive biocatalysts for the conversion of CO_2_ with H_2_ into acetate, as in gas fermentation. Gas fermentation reactors may benefit from biofilm formation, but cell attachment by acetogens is often limited. This study indeed found that the acetogen *Sporomusa ovata* 2663 was mainly planktonic and aimed to increase its biofilm formation through adaptive laboratory evolution. The adaptation strategy consisted of growing *S. ovata* on plastic carriers in bottles with a H_2_:CO_2_ headspace and transferring few carriers to bottles with fresh carriers over eight serial transfers. This procedure resulted in the evolved *S. ovata* 2663-BF, which had a consistent increased propensity to attach. In heterotrophic growth conditions, four times more cells attached to the bottom of well plates in comparison to the wild type. Moreover, twice as many cells adhered to carriers when grown on H_2_:CO_2_. This improved attachment, however, did not lead to higher acetate production rates in simple trickle bed reactors, as the used experimental setup likely stimulated planktonic growth due to a low trickling frequency. Only after some medium replacements to remove planktonic cells, higher gas consumption rates were recorded for the evolved culture. Interestingly, the evolved *S. ovata* had a relevant point mutation in the gene *galU*, encoding UDP-glucose pyrophosphorylase, an enzyme involved in the synthesis of extracellular polysaccharides. Overall, this study demonstrates that cell attachment by *S. ovata* was increased through adaptive laboratory evolution, offering the prospect of investigating the importance of biofilm formation in biofilm-based gas fermentation reactors.

**IMPORTANCE:** Acetogenic bacteria are attractive biocatalysts for the conversion of CO_2_ with H_2_ into acetate, as in gas fermentation and microbial electrosynthesis. Both biotechnologies could benefit from biofilm formation, since biofilm formation retains the catalytic activity inside continuously operated reactors. In addition, biofilm-based gas fermentation reactor systems are advantageous to improve the gas to liquid mass transfer of H_2_ and CO_2_. Moreover, microbial electrosynthesis benefits from biofilms on cathodes to consume H_2_ as soon as it gets generated. Acetogenic bacteria are however often found to only form thin or sparse biofilms. In the current study, we demonstrated that adaptive laboratory evolution is a useful strategy to increase the biofilm formation capabilities of the acetogen, *Sporomusa ovata*. The obtained biofilm improved *S. ovata* is of interest to deepen our fundamental understanding of biofilm formation by acetogens, as well as to assess the importance of biofilm formation for the performance of biofilm-based bioreactors.

## INTRODUCTION

Acetogenic bacteria are attractive biocatalysts for the development of CO_2_ utilization and energy conversion technologies. Acetogens use CO_2_ as a terminal electron acceptor with H_2_ as an electron donor for energy conservation via the Wood-Ljungdahl pathway, leading to the formation of mainly acetate (1, 2). The H_2_ can be produced from renewable electricity by electrolysis and brought together with CO_2_ in an acetogenic bioreactor, as in gas fermentation applications (3). Alternatively, an electrode (cathode) can deliver H_2_ directly into a bioreactor, as in microbial electrosynthesis (MES) (4). Several studies have described the acetogenic bacterium *Sporomusa ovata* as a highly interesting biocatalyst for both type of applications, due to its high acetate production rates (5–7).

Several reactor systems for gas fermentation are biofilm-based. For instance, in trickle bed reactors for gas fermentation, H_2_ and CO_2_ are converted by microorganisms attached to carrier materials present in the gas phase, while a liquid phase trickles over the carriers to provide nutrients (8, 9). Also, membrane biofilm reactors are used for gas fermentation and are based on biofilm formation of microorganisms on a membrane, through which gaseous components diffuse (8, 10). Similarly, in microbial electrosynthesis, microorganisms often attach to the cathode (5, 11), which allows the microorganisms to consume H_2_ as soon as it is generated by the electrode. Biofilm formation in these systems has the additional advantage that the microorganisms and their catalytic activity remain contained in the bioreactor, while their products can be harvested from the liquid phase by continuous operation. Excessive biofilm formation, on the other hand, can negatively impact the activity of cells in a biofilm, as the supply of substrates and the removal of products in thick biofilms gets limited by diffusion (8). Investigations of the role of biofilm formation in those bioreactors thus forms an essential step towards the further development of CO_2_ utilization biotechnologies.

Several studies have already reported biofilm formation by acetogenic bacteria, including *S. ovata*, on various substrates (**Table S1**). Attachment of pure cultures of acetogens was demonstrated in trickle bed and membrane biofilm reactors, as well as on cathodes in MES bioreactors (**Table S1**). Interestingly, *Sporomusa* was identified as sole acetogen in methanogenic biofilms grown on H_2_ and CO_2_ (12). Despite these findings, only few studies have specifically placed biofilm formation by acetogens in focus (13), while the molecular mechanisms underlying their attachment remain completely uncharacterized. Furthermore, most existing studies only reported thin biofilms and sparse attachment, leading to incomplete surface coverage (**Table S1**). Enhancing biofilm formation by acetogens could therefore offer valuable improvements for biofilm-based production biotechnologies. However, acetogenesis from H_2_ and CO_2_ yields very little energy, making acetogens balance at the so-called ‘thermodynamic limit of life’ (2). It remains unclear whether acetogens can allocate sufficient energy for the production and secretion of extracellular polymeric substances (EPS) (i.e. polysaccharides, proteins or nucleic acids) (14), which are essential for surface attachment and biofilm matrix formation.

One commonly used strategy to improve a not well-understood phenotypic trait of a microorganism is Adaptive Laboratory Evolution (ALE). By exposing microorganisms to a selection pressure over a prolonged period of time, beneficial (random) mutations are enriched and better adapted microorganisms are obtained (15–17). ALE has already been applied to various acetogenic bacteria to improve or establish desired characteristics (18–20). For instance, ALE was used to increase the tolerance to O_2_ (21) and methanol of *S. ovata* (22). ALE has also been successful in increasing biofilm formation of aerobic microorganisms. For example, ALE enhanced the cathodic autotrophic biofilm growth capabilities of the aerobe *Kyrpidia spormanii* (23). In addition, ALE increased biofilm production of the heterotrophic aerobe *Burkholderia cenocepacia* (24). The latter study selected for biofilm formation by repeated serial transfers of plastic beads (with adhered cells) to fresh medium with new clean beads. So far, ALE has not been tested to increase biofilm formation of anaerobic bacteria, such as acetogens, while overall the role of acetogenic biofilms in biofilm-based bioreactors remains not well understood.

This study aimed to investigate the native biofilm formation properties of biotechnologically relevant acetogen *S. ovata* strain 2663, as well as to increase its attachment and biofilm formation capabilities through ALE. Hereto, *S. ovata* was grown in bottles filled with carriers incompletely covered with liquid medium in an atmosphere containing H_2_ and CO_2_ (resembling trickle bed reactors). ALE was performed through serial transfers of few of the carriers to bottles with fresh carriers. Attachment, planktonic growth, as well as growth yields, of the resulting biofilm-increased *S. ovata* 2663-BF were compared with those of the wild type (WT) *S. ovata* in both heterotrophic and autotrophic growth conditions. Furthermore, the performance of *S. ovata* 2663-BF was compared with that of the wild type in simple trickle bed reactors as an attempt to investigate the importance of biofilm formation in such reactor systems. Finally, insights into the molecular basis of the altered phenotype were obtained through genome comparison.

## RESULTS

### Trickle bed experiments with wild type *S. ovata*

Initial experiments evaluated the performance of WT *S. ovata* 2663 in simple trickle bed reactors and assessed its intrinsic capacity to attach. Simple trickle bed reactors were bottles filled with plastic carriers and a small amount of medium in a H_2_:CO_2_ containing headspace. In a first experiment, gas consumption by *S. ovata* was monitored in bottles with batch gas supply and with or without carriers (manually turned around daily, otherwise static incubation). In the abiotic controls, a gradual drop in pressure was observed, illustrating the loss of gas due to repeated pressure measurements (**Fig. 1A**). Both biotic treatments showed a faster drop in the pressure than the abiotic controls, showing the gas consumption activity of *S. ovata*. Interestingly, in the bottles with carriers, the pressure drop was at least ten times faster than without carriers (**Fig. 1A**). This relates to the ten times larger surface area of the gas-liquid interface (**Supplementary Calculation**), demonstrating that the carriers made the bottles function as trickle bed reactors.

**FIG 1.**
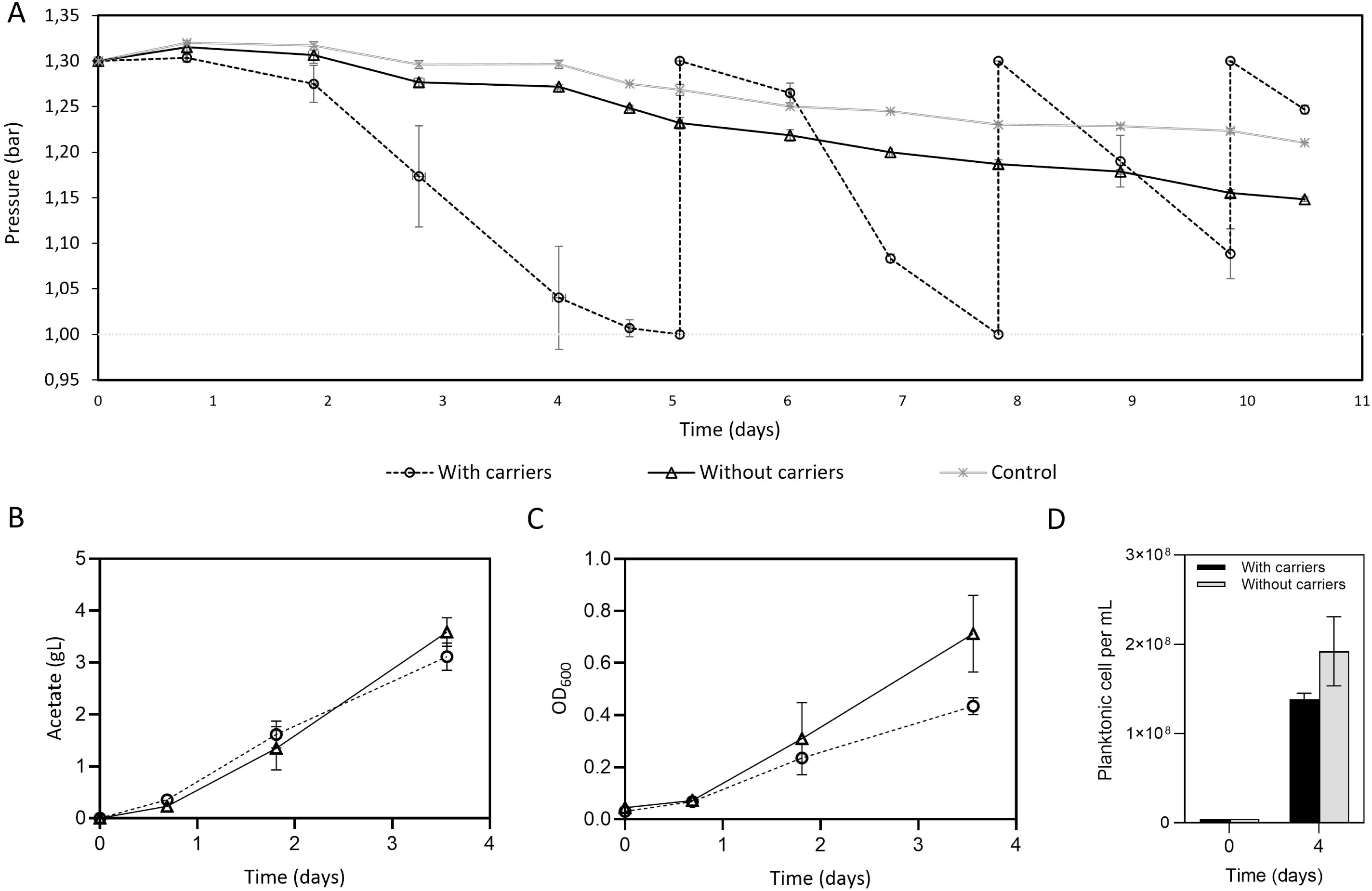
Performance of *S. ovata* DSM-2663 (WT) under autotrophic growth conditions with and without the presence of carriers. All bottles were statically incubated and manually turned around at least daily. **(A)** Gas (H_2_:CO_2_) consumption (measured as a decrease in overpressure) by *S. ovata* DSM-2663 in bottles with or without carriers and in abiotic control bottles. At the start, these bottles contained 1 bar of N_2_:CO_2_ (80:20) gas (represented by the dashed line in the bottom) and 0.3 bar overpressure of H_2_:CO_2_ (80:20) gas. For the bottles with carriers, H_2_:CO_2_ (80:20) gas was reinjected on day 5, 8 and 10, when the overpressure approached or reached zero. (**B**) (**C**) and (**D**) Increase of respectively the acetate concentration, optical density (OD_600_) and planktonic cell numbers by *S. ovata* DSM-2663 in bottles with or without carriers. In these bottles, H_2_:CO_2_ (67:33) gas was continuously supplied to maintain a constant overpressure of 1.5 bar. All points represent the average of at least triplicated bottles, while the error bars show the standard deviation.

In a second experiment, acetate concentrations and planktonic growth were monitored in reactors with continuous H_2_:CO_2_ gas supply and with or without carriers. This condition led to similar acetate concentrations with and without carriers after four days of incubation (**Fig. 1B**), showing that carriers did not have the same advantage, as observed under batch gas supply (**Fig. 1A**). Interestingly, however, in the bottles with carriers, the optical density (OD_600_) of the liquid phase was about 40% lower than without carriers after four days of incubation (**Fig. 1C**), while planktonic cell densities were about 30% lower (significant difference, p < 0.05) (**Fig. 1D**). A significant number of cells had thus likely attached to the carriers, as similar overall cell numbers could be expected from the comparable acetate concentrations (**Fig. 1B**). This suggests that the WT *S. ovata* 2663 has the capacity to attach to surfaces and does not stay purely planktonic in autotrophic growth conditions.

### ALE towards increased biofilm formation

Starting from one of the bottles with carriers of the first trickle bed experiment (**Fig. 1A**), ALE was carried out by repeatedly transferring few carriers to new trickle bed reactors with fresh medium and fresh carriers (**Fig. 2A**). In this way, only cells associated to the carriers were transferred, creating a selection pressure for attachment. Over the course of the ALE experiment, the time from the transfer until detectable gas consumption decreased. For the first transfers, the time between the transfer and the consumption of 0.3 bar of gas overpressure was about 16 to 23 days, while for the last transfers, this time was about 6 to 8 days. As a consequence, the intervals between the serial transfers became shorter during the course of the ALE experiment (**Fig. S1**). At the later transfers, the culture was also propagated in heterotrophic growth medium to allow the preparation of frozen stocks. When grown under heterotrophic growth conditions, the evolved *S. ovata* 2663 culture obtained after eight serial transfers, visibly differed from the WT *S. ovata* (**Fig. 2B**). The WT *S. ovata* homogenous grew throughout the liquid phase and was visibly planktonic, in contrast to the evolved *S. ovata*, which formed a visible biofilm on the sides of glass tubes. As this was the desired phenotype, ALE was ended at this timepoint. The adapted culture with increased biofilm features obtained at the end of the ALE procedure is further called *Sporomusa ovata* 2663-BF (or BF culture in short).

**FIG 2.**
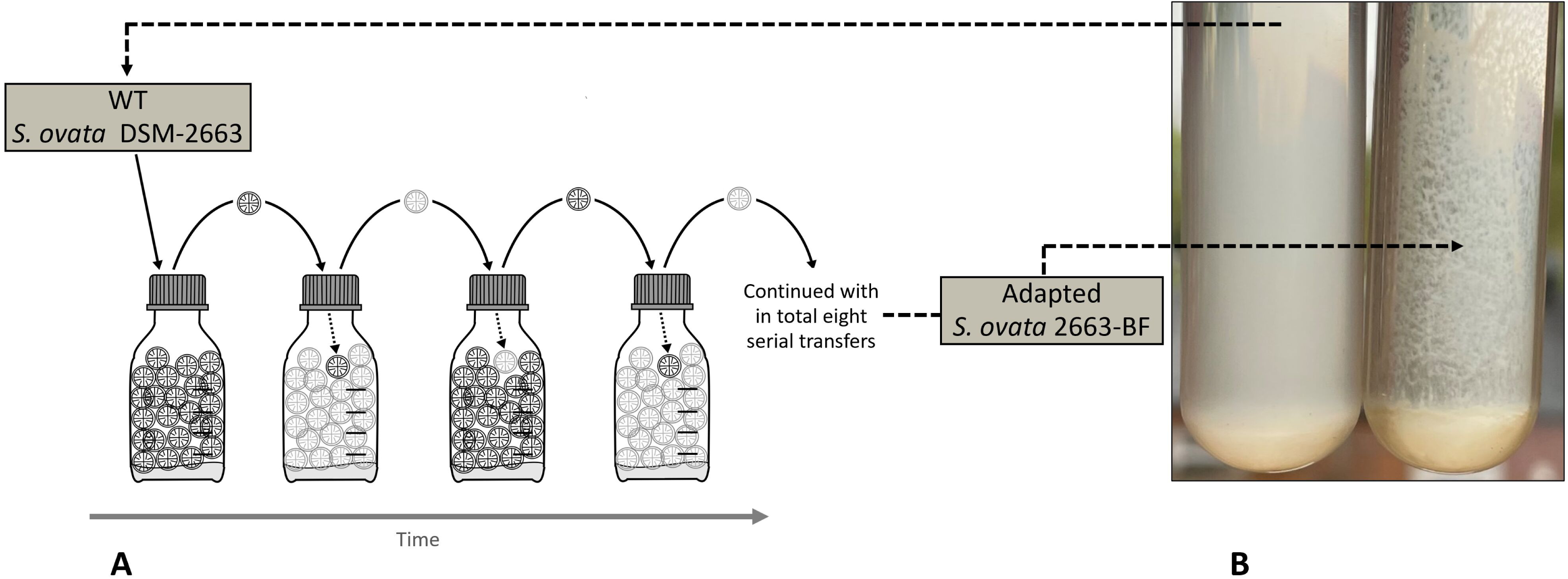
(**A**) Conceptual scheme illustrating the adaptive laboratory evolution strategy used to increase biofilm formation by *S. ovata.* The WT *S. ovata* DSM-2663 was grown in simple trickle bed reactors under an atmosphere of N_2_, CO_2_ and H_2_ (batch gas supply, manual turned around several times per week). When the gas at overpressure was consumed, H_2_:CO_2_ gas was reinjected. After two to three reinjections, two carriers were transferred to a new reactor with fresh medium and fresh carriers. The change in the grey colour of the carriers in the scheme illustrates the transfer between different bottles. Eight transfers were performed to obtain the final adapted *S. ovata* 2663-BF with increased biofilm formation properties. (**B**) Picture showing the visual difference between the WT *S. ovata* and the adapted BF culture, when grown in glass Hungate tubes under heterotrophic growth conditions. The WT was mainly planktonic, while the BF culture was rather associated to the side of the tube.

### Comparison of biofilm formation in heterotrophic growth conditions

Heterotrophic growth conditions were chosen to compare the biofilm formation characteristics between the WT and the BF *S. ovata*, since this allowed growth in well plates in an anaerobic chamber. After three days of growth in well plates, the OD_600_ of the planktonic phase, as a measurement of planktonic growth, was about four times lower for the BF culture compared to WT (**Fig. 3A, left**). Conversely, the biofilm formation of the evolved *S. ovata* was about ten times higher than the wild type, as quantified by a crystal violet assay (**Fig. 3A, right & 3B**). In addition, the increased biofilm formation by the evolved *S. ovata* corresponded with an at least four times higher cell numbers attached to the well plates than with the WT (**Fig. 3C**), while in the planktonic phase of the BF culture had seven times less cells than for the WT (**Fig 3C**). Similar experiments using the crystal violet assay were further repeated to acquire a time resolution, in which the same trends were reproduced and biofilm formation was found to be stable over at least a few days (quantified from day 2-4) (**Fig. 3D**). In all, these results demonstrate that the ALE procedure increased the capacity of *S. ovata* to attach to surfaces and form biofilms, at least under heterotrophic growth conditions.

**FIG 3.**
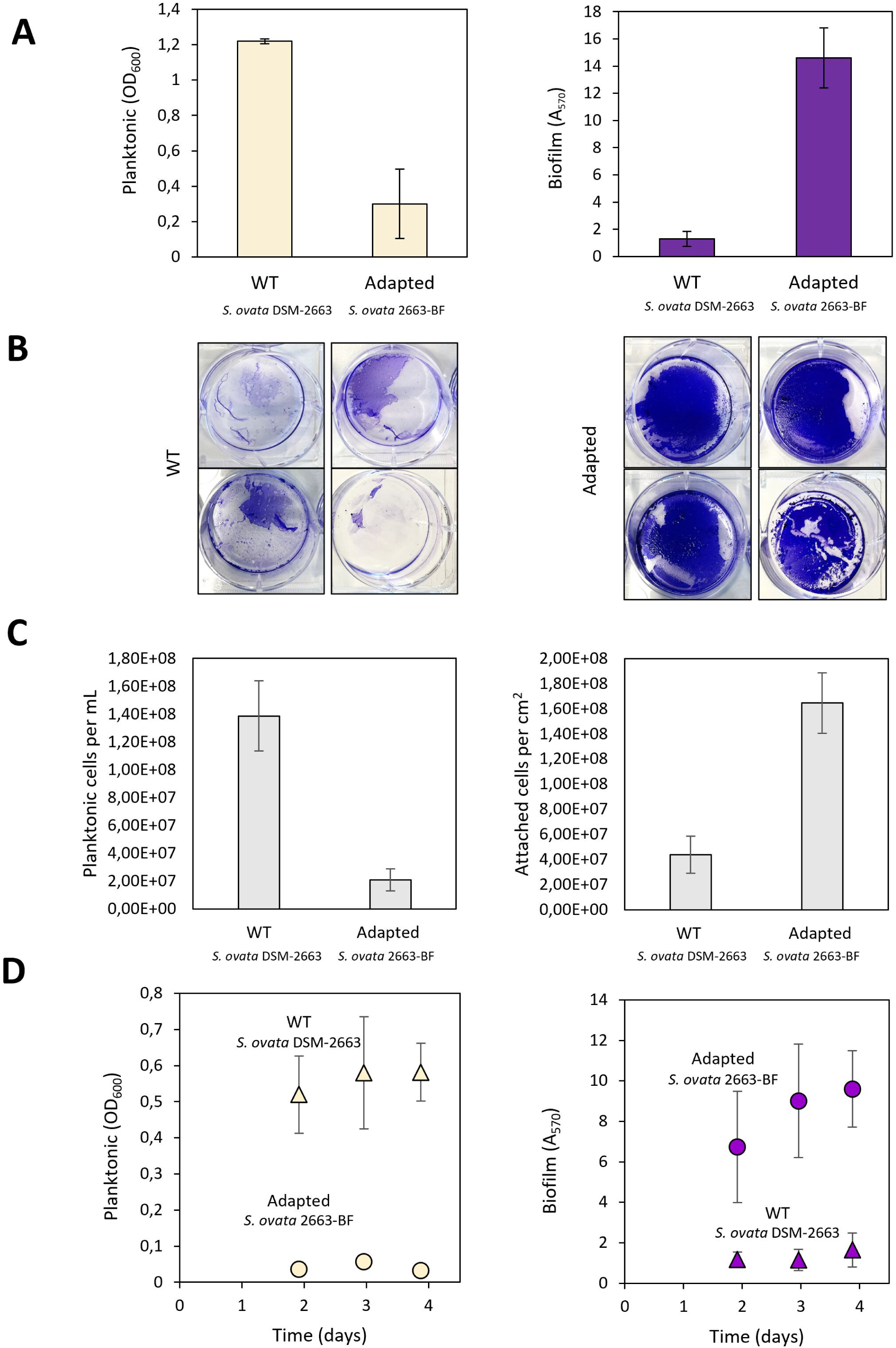
Comparison of planktonic growth and biofilm formation by the WT *S. ovata* DSM-2663 versus the adapted *S. ovata* 2663-BF in heterotrophic growth conditions and when grown in well plates. (**A**) Planktonic growth was assessed by the optical density (OD_600_) (left) and biofilm formation was quantified by the crystal violet assay (A_570_) (right) after 3 days of incubation. (**B**) Representative pictures of biofilms stained with crystal violet before extraction with methanol from different repeated runs, all after 3 days of incubation, with WT (left) and BF (right). (**C**) Cell density of the planktonic phase (left) and the number of cells per cm^2^ in the biofilm attached to the bottom of the well (right) as measured using IFC after 3 days of incubation. (**D**) Planktonic growth measured by the optical density (OD_600_) (left) and biofilm formation quantified by the crystal violet assay (A_570_) (right) at three different time points. The height of all bars or points represents the average of measurements of three to nine wells, while the error bars show the standard deviation.

### Comparison of performance in trickle bed reactors

The performance of the BF *S. ovata* was further compared with that of the WT in simple trickle bed reactors using continuous gas supply. Medium was replaced every few days to remove planktonic cells and stimulate biofilm growth. Even though both cultures were inoculated with equal cell numbers, the WT started to accumulate acetate faster, while a lag phase of two days was observed in the acetate production by the BF culture (**Fig. 4A**). After four days, the acetate concentration was double so high for WT (3.1 ± 0.3 g·L^-1^) than for BF (1.4 ± 0.7 g·L^-1^), which was also reflected by a lower pH and higher OD_600_ for WT than for BF (**Fig. 4B and S2**). Ethanol concentrations remained under the detection limit throughout the entire experiment.

**FIG 4.**
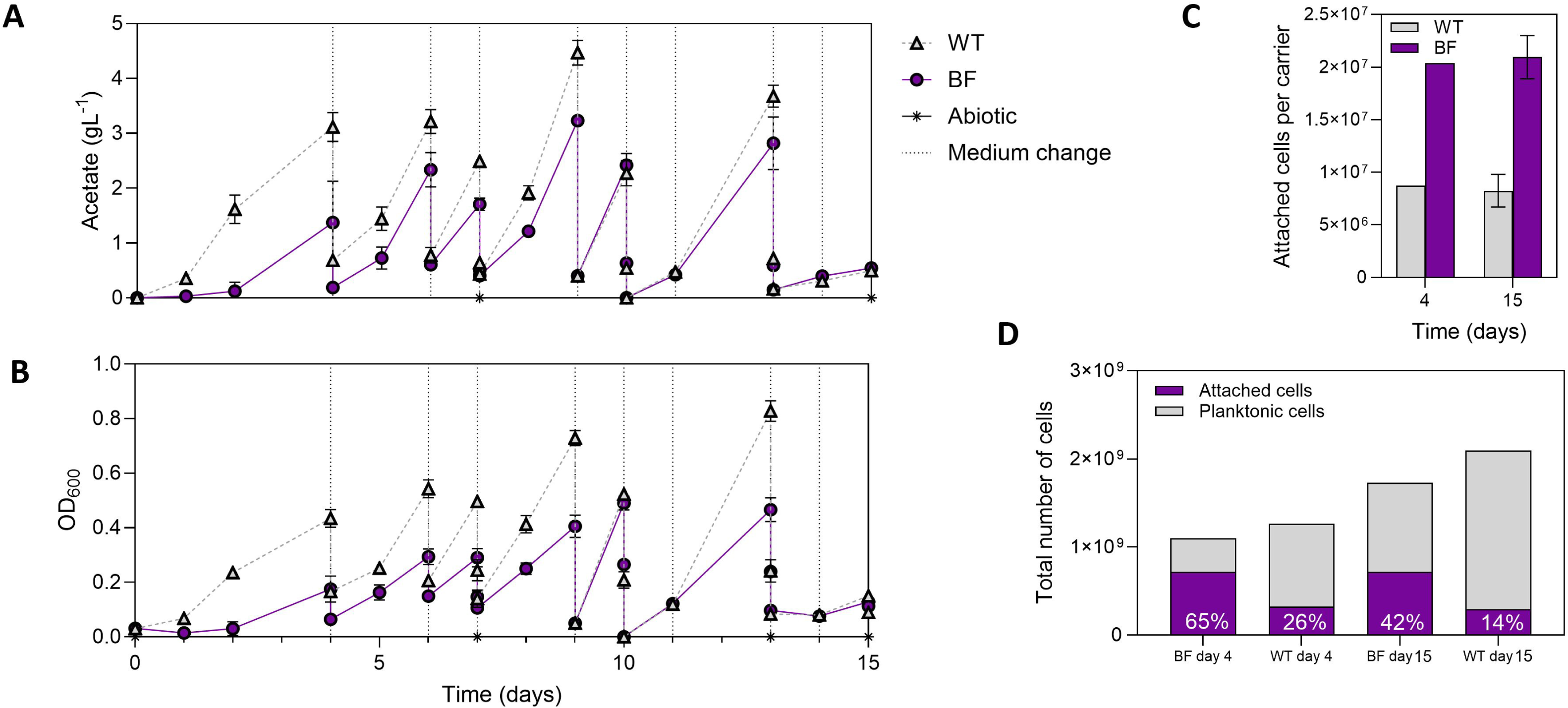
Comparison of the WT *S. ovata* DSM-2663 and the adapted *S. ovata* 2663-BF in autotrophic growth conditions when grown in simple trickle bed reactors. In all bottles, H_2_:CO_2_ (4:2) gas was continuously supplied to maintain a constant overpressure of 1.5 bar. Bottles were manually turned around once per day to create trickling effect, otherwise static incubation. Medium was replaced to stimulate biofilm growth, as indicated by the vertical lines. (**A**) and (**B**): Change of respectively acetate concentrations and optical densities (OD_600_) over time. (**C**) The number of attached cells per carrier after 4 and 15 days, as measured using IFC. (**D**) The total number of attached and planktonic cells at day 4 and 15, as calculated from the number of attached cells per carrier (Fig 4C) with 36 carriers and the OD_600_ at day 4 and 13 (before medium replacement) (Fig 4B) with 10 mL liquid phase. Data points represent the average of at least three replicates, while the error bars show the standard deviation, with the exception of the number of attached cells on the carriers at day 4, for which only one replicate bottle was measured.

At day four, one of the four replicated bottles of each culture was harvested to quantify the cells attached to the carriers. The BF strain had 2.3 times more cells attached (2.0·10^7^ cells·per carrier) than the WT (9·10^6^ cells per carrier) (**Fig 4C**), even though it had produced less acetate, and therefore likely had grown less extensively. It was further calculated that 67% of the BF cells were attached, while this was only 26% for WT cells at day 4 (**Fig 4D**). This confirms that also in autotrophic growth conditions, the BF strain has an increased tendency to attach and form biofilms in comparison to the WT strain.

Medium was replaced at day 4 in the remaining three bottles of each culture, which was repeated every few days over a period of 10 days. After the medium replacements, the OD_600_ was not completely reset to zero (**Fig. 4B**), which indicates that not all planktonic cells were removed, or that some attached cells got suspended by the addition of fresh medium. During the 10-day period, the acetate production rate of both strains strongly increased (**Fig. 4A**), suggesting increasing cell numbers in the reactors of both cultures. Moreover, the BF and WT culture reached comparable acetate rates, up to 2.27 ± 0.19 g·L^-1^·day^-1^ for WT and 2.42 ± 0.17 g·L^-1^·day^-1^ for BF after the fourth medium replacement (day 9), but in general the acetate titers remained slightly lower for BF culture than for WT (e.g. 3.23 ± 0.08 g·L^-1^ for BF versus 4.47 ± 0.19 g·L^-1^ for WT at day 9) (**Fig. 4A**). Accordingly, the pH dropped less in between the medium replacements for BF than for WT (e.g. pH 5.71 for BF vs 5.32 for WT at day 9) (**Fig. S2**). For both strains, the OD_600_ increased between medium replacements (**Fig 4B**), indicating that at least a part of the growth occurred planktonically, or otherwise resulted from the suspension of biofilm grown cells. The OD_600_ increase between medium replacements was less pronounced for the BF culture than for WT (OD_600_ of 0.405 for BF vs OD_600_ of 0.729 for WT at day 9), which might be due to a higher number of attached BF cells (**Fig. 4C**). Overall, the acetate and OD_600_ measurements indicate that the BF culture performed comparable to the WT in a system in which planktonic growth apparently dominated.

In order to assess the activity of the attached cells (and any remaining planktonic cells), the continuous gas supply was halted after some of the medium replacements for a few hours to enable measurement of the gas consumption rates (i.e. decrease of pressure per time unit). At all instances, the pressure dropped much faster in the biological treatments than in the abiotic controls (**Fig 5A** and **S3**), confirming the activity of both cultures. Moreover, at two instances, the BF culture had significantly higher gas consumption rate than the WT (**Fig 5A**). At day 4, the BF gas consumption rate was double as high as that of WT, which is possibly due to its higher cell numbers attached to the carriers (**Fig 4C**), but could relate to the WT cells having to overcome stationary phase, which this culture likely had reached before the medium transfer (pH < 5.5 at day 4 for WT) (**Fig S2**). At day 9, the gas consumption rate of the BF culture (18.83 mbar·h^-1^) was also double so high as that of WT (9.26 mbar·h^-1^) (**Fig 5**), which likely reflects the higher number of attached BF cells, as both strains had a highly comparable performance in terms of acetate production rate and OD_600_ increase before the medium replacement (**Fig 4A &B**). Gas consumption rates were also measured at other time points (day 6 and 11), but were then not significantly different between the two cultures (**Fig 5A**).

**FIG 5.**
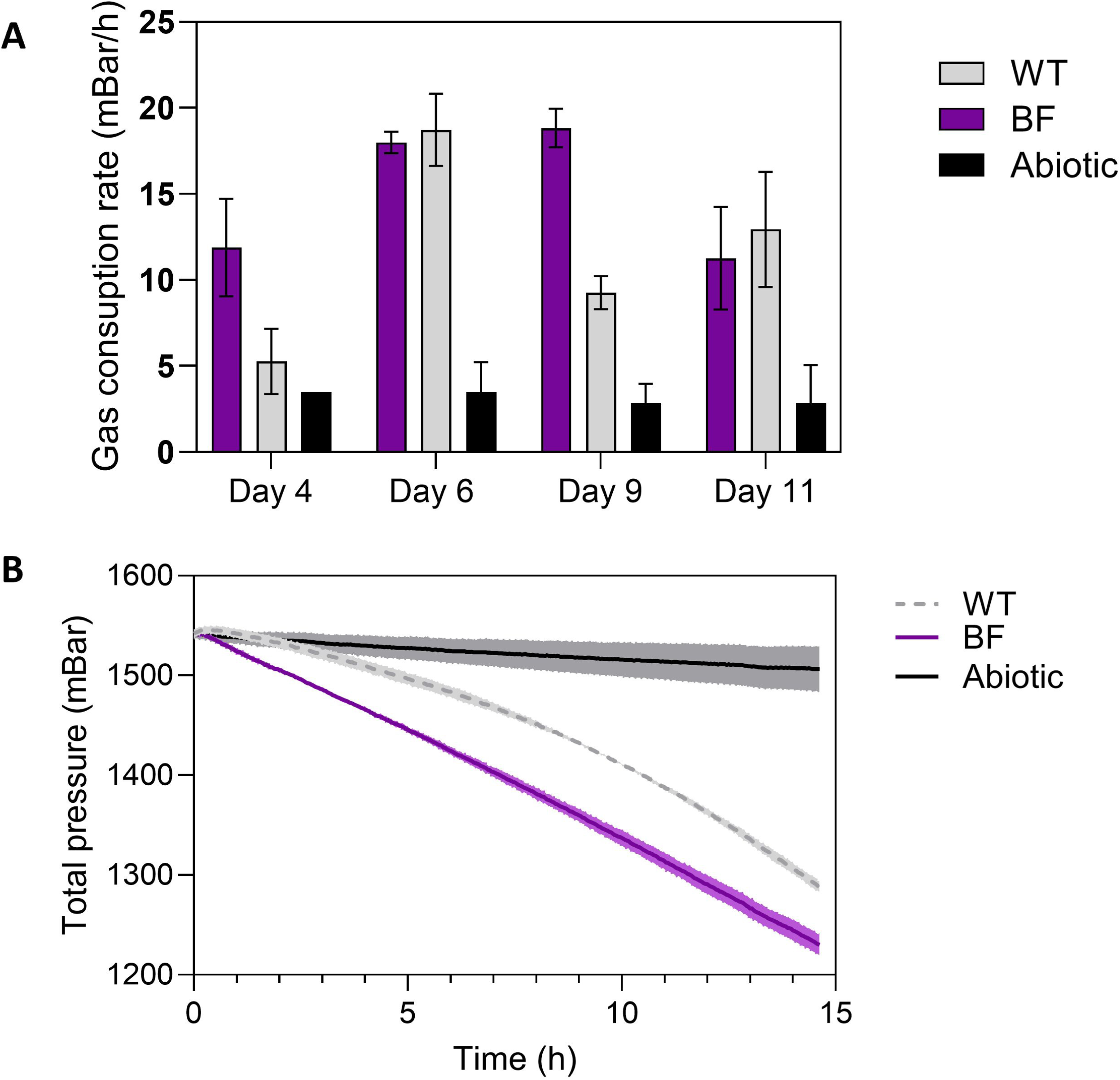
(**A**) Gas consumption rates of the WT *S. ovata* DSM-2663 and the adapted *S. ovata* 2663-BF at four different time points, when grown in autotrophic growth conditions in simple trickle bed reactors. To measure these gas consumption rates, the continuous gas supply was halted after the medium replacements. All gas consumption rates were calculated from the pressure decrease during the second hour (first hour as equilibration time). The height of the bars shows the average of triplicate bottles, while the error bars reflect the standard deviation. (**B**) Pressure decrease measurement for a period of 15 hours during day 9, comparing gas consumption by the WT and BF culture, as well as in abiotic conditions. The darkest line represents the average of three replicate bottles, while the shaded area shows the standard deviation.

At some instances the pressure measurements were continued overnight (up to 16 hours). Interestingly, around day 9, this resulted in an almost constant gas consumption rate (linear pressure drop) for the BF culture, while the gas consumption rate of the WT increased over time (exponentially decreasing pressure) (**Fig. 5B**). This difference in the shape of the curves was also recorded after several of the other medium replacements (**Fig. S3**). This observation suggests that gas consumption by the WT depended on (exponential) growth, while less growth occurred for the BF culture, possibly because more cells remained in the reactors after the medium replacement, thanks to its higher attachment.

At the end of the experiment (after 15 days), all remaining bottles were harvested to quantify the cells attached to the carriers. The adapted *S. ovata* had still about 2.53 times more attached cells (2.09·10^7^ cells per carriers) than the WT *S. ovata* (8.2·10^6^ cells per carrier) (**Fig 4C**). Using the OD_600_ values measured before the medium replacement at day 13 (growth was not observed anymore between day 13 and 15), the number of planktonic cells was estimated and compared with the number of attached cells at day 15. These calculations showed that about 42% of the BF cells were attached, while this was 14% for WT cells at the end of the experiment (**Fig 4D**). The relative number of attached cells thus decreased over the course of the experiment (**Fig 4D**), while the BF *S. ovata* was persistent in its increased attachment in comparison to WT.

### Comparison of growth yields

The growth yield of the BF and WT *S. ovata* was determined both under heterotrophic and autotrophic growth conditions (**Fig. 6**). Growth yields, incorporating both the increase of attached and planktonic cells, were not significantly different between the two cultures, both when expressed as the increase of cell numbers or as cell dry weight per mole of acetate produced (p > 0.05), indicating that the increased propensity of the BF culture to form biofilms did not impair its capacity to grow.

**FIG 6:**
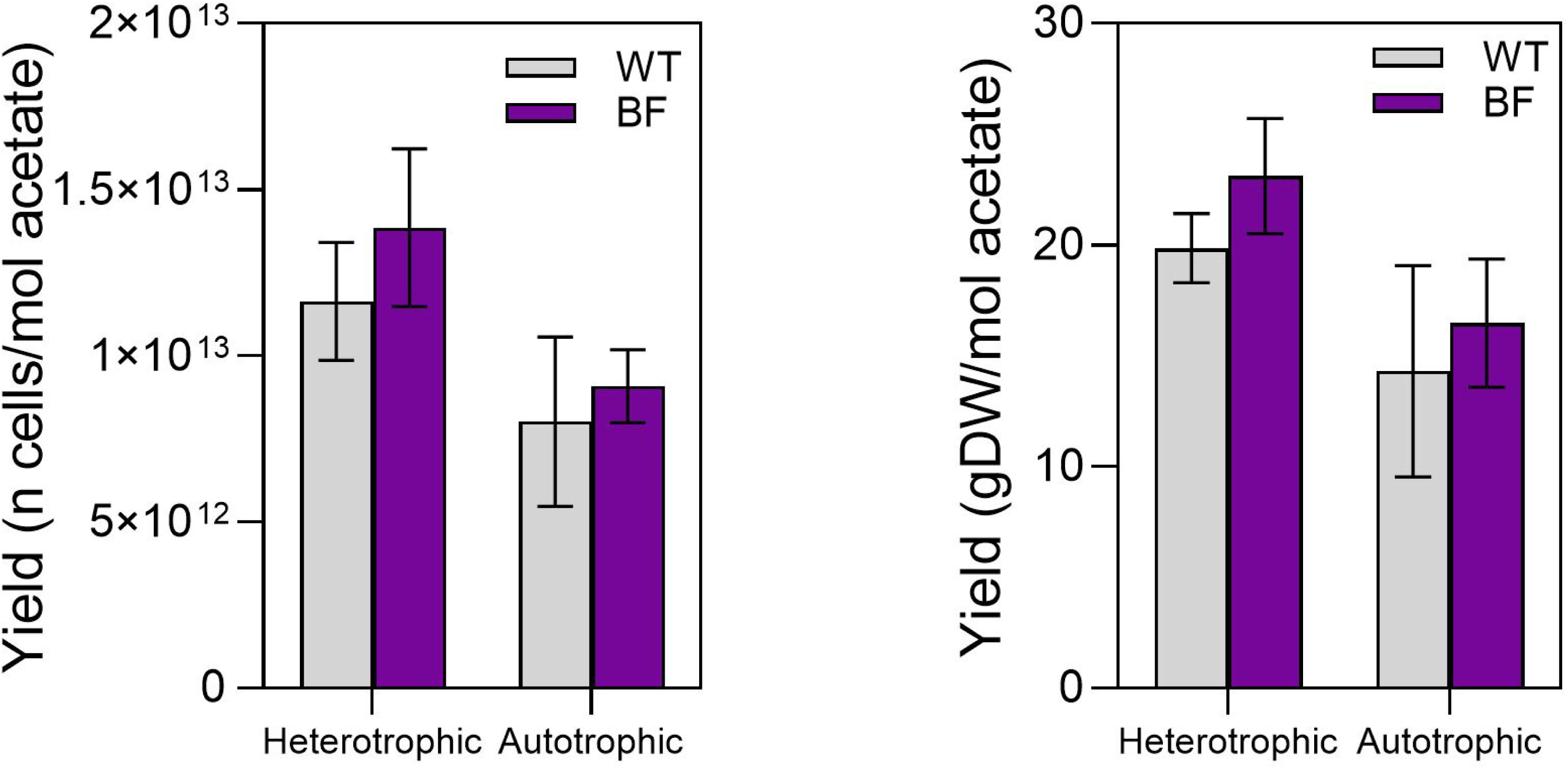
Growth yields for WT *S. ovata* DSM-2663 and the adapted *S. ovata* 2663-BF both in heterotrophic and autotrophic growth conditions, expressed as either (**A**) increase of cell numbers per mol of acetate produced; or (**B**) increase of cell dry weight (g) per mole of acetate produced. Both attached and planktonic cells were included in the growth yields. The height of the bars shows the average of five replicated bottles, while the error bars reflect the standard deviation.

### Genomic comparison

To obtain insights into the biofilm formation mechanisms of the evolved *S. ovata* 2663-BF, we performed a genome analysis comparing the genome of the WT and BF culture. Mapping the sequencing reads to the reference genome indicated several mutations in the BF versus the WT *S. ovata* 2663 (**Table S2**). Overall, 11 mutations were found in protein-coding genes, while there were eight mutations in intergenic regions, and two mutations in ribosomal RNA genes. Of the eight mutations in intergenic regions, five were adjacent to long homopolymer stretches and thus likely the result of sequencing errors (**Table S2**). The three remaining intergenic mutations were not upstream of any coding sequences and thus unlikely to affect the bacterium’s physiology. Of the 11 apparent mutations in protein-coding genes, nine were found in genes with multiple copies (*cdhC* and phage tail protein) and likely result from inaccurate mapping of reads between the different copies (**Table S2**). The two remaining mutations in protein-coding sequences included a frameshift mutation in hypothetical protein G571_RS0122020, of which the role cannot be determined; as well as most interestingly a missense variant in *galU* (**Table S2**). This single-nucleotide mutation at position 434 in the *galU* gene changed the encoded amino acid from a leucine to a histidine (L145H). The gene *galU* encodes the enzyme UTP-glucose-1-phosphate uridylyltransferase (UGPase) (**Fig. 7**), which catalyses the formation of UDP-glucose from glucose-1-phosphate and UTP and has previously been related to the synthesis of extracellular polysaccharides (25, 26). This mutation was found in 89 of the 90 sequencing reads mapped to this gene (**Table S2**), showing that the mutation had become almost completely established in the BF culture.

**FIG 7.**
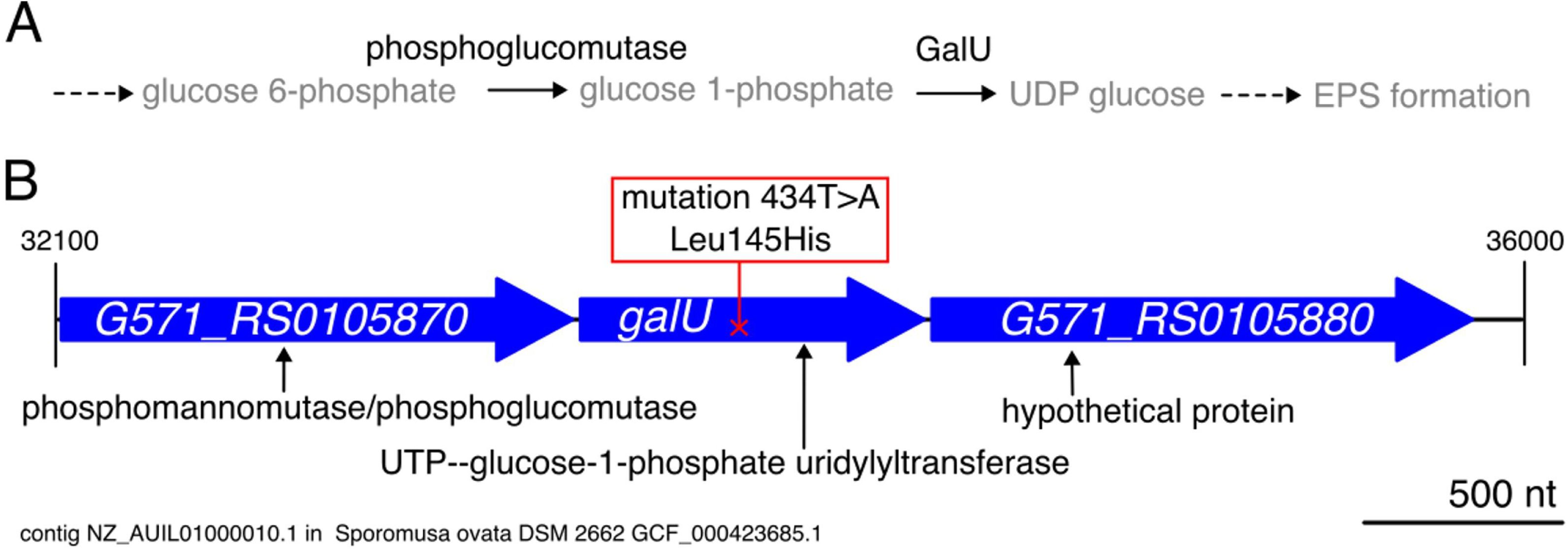
Schematic representation of (A) a pathway for EPS production in which UTP-glucose-1-phosphateuridylyltransferase (UGPase) is expected to be involved and (B) the location of the mutation in the *galU* gene causing an altered amino acid sequence of the UGPase enzyme.

### Structural comparison of the UGPase of WT and BF *S.ovata*

We investigated whether the point mutation in *galU* could have affected the enzyme activity of the encoded UGPase, through the generation of structural models of WT and the mutant enzyme (L145H). In UGPase enzymes, glucose-1-phosphate makes a nucleophilic attack on the α-phosphate of UTP to form UDP-glucose, while magnesium ion (Mg^2+^) at the active-site plays a key role in the catalysis by charge neutralization and by positioning the substrates (27–32). According to our comparative structure analysis using AlphaFold and AlphaFill, the mutated residue (L145H) is far from the active-site pocket, with over 16 Å distance to Mg^2+^ (**Fig. 8A**). However, the imidazole side chain of H145 in the mutant is within interaction distance to the backbone carbonyl oxygen of a tyrosine 210 (Y210) (4.2 Å distance, **Fig. 8A**). Y210 is highly conserved among UGPase orthologs and structural studies assigned Y210 as one of the residues forming a hydrophobic cap at the base of the glucose ring in the active-site (27). Moreover, in our structural model of the *S. ovata* UGPase, the hydroxy group of Y210 is within 4.4 – 6.0 Å distances from the oxygen atoms of the glucose moiety of UDP-glucose, indicating a role for Y210 in binding and/or catalysis. Thus, interaction between H145 and Y210 can possibly lead to higher catalytic efficiency in the mutant by affecting substrate binding or through better positioning of the substrate(s) for catalysis. In the WT enzyme, L145 is not able to make this interaction with Y210 due to its inert side chain.

**FIG 8.**
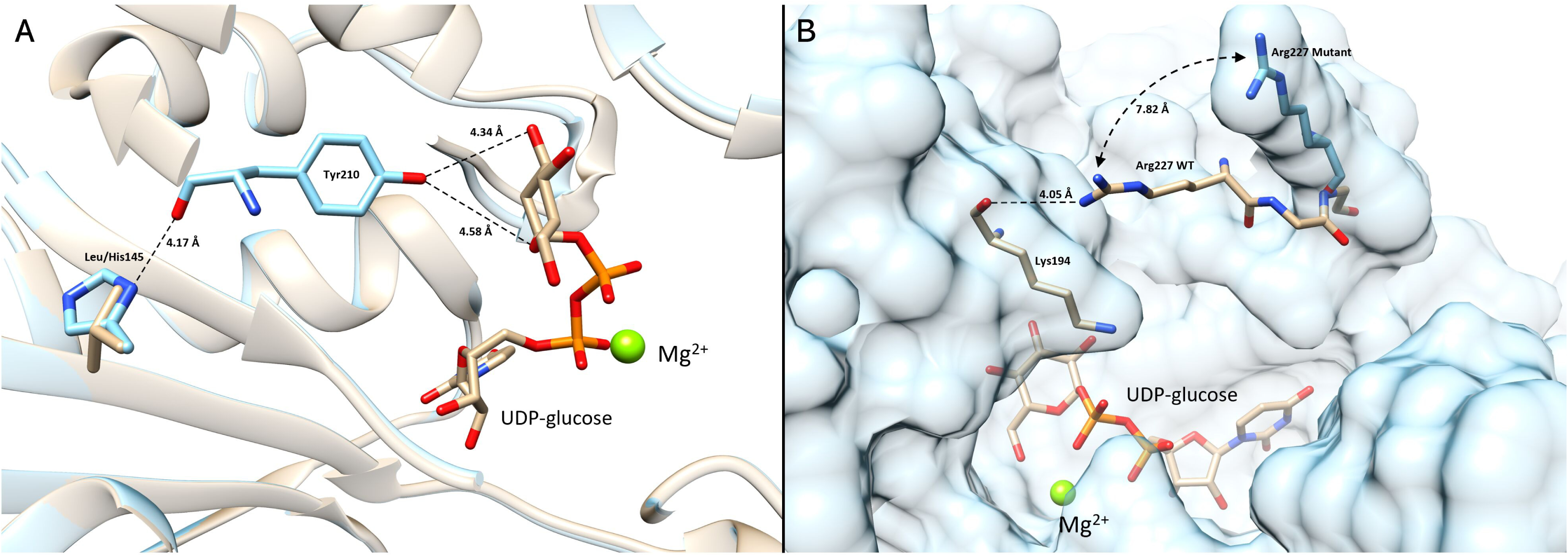
(A) Overlaid active-site of WT and mutant *S.ovata* UGPase highlighting the mutation (L145H) site and possible interactions between His145 side chain and Tyr210. The brown structure is WT UGPase and the blue structure is the mutant. The distances between Tyr210 side chain and glucose moiety of UDP-glucose are shown. The green sphere is the magnesium ion. (B) Overlaid structures of WT and mutant *S.ovata* UGPase. Brown structure is WT enzyme and blue structure is the mutant. The surface representation is only shown for mutant enzyme for clarity. Highlighted are the Arg227 residue that is positioned differently between the two models and the likely interaction with Lys194 in the WT enzyme.

In addition, structural comparison of WT vs. L145H mutant UGPase revealed a significant difference in the position of Arginine 227 (R227) (**Fig. 8B**), which resides on a mobile gatekeeping loop at the entry of an access tunnel to the active site pocket, according to earlier studies (27) and our analysis by Caver Web software. In the WT enzyme, R227 is 7.8 Å closer to the active-site and partially closes the access tunnel, with its side chain possibly interacting with the backbone of two active-site residues, Lysine 194 (K194) and Glutamate 193 (E193) (27) (**Fig. 8B**). In the L145H mutant, these interactions are absent. Although AlphaFold 3 models of the enzymes had a higher degree of uncertainty in the loop region including R227, such differences in the mutant possibly affect the size, access and organization of active-site pocket and/or conformational changes during catalysis or binding events. It has been well documented that various conformational changes and active-site dynamics play a role in the bi-substrate mechanism of UGPases (28).

Additionally, it is possible that the L145H mutation favorably affected the stability of the enzyme, increasing overall catalytic performance. In short, our structure analyses show that the point mutation in *galU* most probably increased the activity of UGPase enzyme, thereby leading to increased EPS formation and cell attachment.

## DISCUSSION

### ALE increased attachment by *S. ovata* 2663

Inherent biofilm formation of acetogens, such as *S. ovata*, is often considered thin or with sparse attachment of cells (**Table S1**). Throughout this study, we found evidence that WT *S. ovata* 2663 was capable of attaching to solid surfaces to a certain extent. In heterotrophic growth conditions, some attachment to the bottom of well plates was observed (**Fig. 3**), while some WT cells were also found to attach to the plastic carriers of the trickle bed reactors under autotrophic growth conditions (**Fig. 1 and 4**). In one MES study, *S. ovata* 2663 was found to have the most pronounced cathodic biofilm formation of various *Sporomusa* strains (**Table S1**) (33). Nevertheless, it should be noted that the WT *S. ovata* 2663 mainly grew planktonic both in auto- and heterotrophic growth conditions, as was for instance visually observed from growth in Hungate tubes (**Fig. 2**), while at least about 75% of the WT cells remained planktonic when grown in trickle bed reactors (**Fig. 1D and 4D**).

To increase attachment and biofilm formation by *S. ovata* 2663, we performed an evolution experiment. Our ALE strategy consisted of transferring carriers with attached cells to fresh trickle bed reactors over a period of six months. During these serial transfers (eight in total), H_2_:CO_2_ gas consumption became faster (**Fig. S1**), suggesting that increasingly more biomass adhered to the carriers, and thus that *S. ovata* attachment was enhanced over the transfer series. Besides the selection pressure for attachment, also the favourable gas to liquid mass transfer conditions on the carriers (improved access to the gaseous substrates H_2_ and CO_2_) (**Fig 1A**) likely contributed to in the adaptation towards increased attachment and biofilm formation.

The resulting adapted *S. ovata* 2663-BF culture had significant increased attachment and biofilm formation properties. In heterotrophic growth conditions, the BF culture had increased biomass and higher cell numbers attached to well plates, while its planktonic growth was strongly reduced in comparison to the ancestral *S. ovata* strain (**Fig. 3**). In addition, in autotrophic growth conditions, about double so many BF than WT cells attached to the carriers (**Fig. 4B**), whereas also the relative number of attached cells was highest for the BF *S. ovata* throughout the whole trickle bed experiment (**Fig. 4D**). Moreover, the pressure consumption curves (**Fig. 5B and S3**) also suggested the increased attachment of the BF culture, as the shapes of these curves indicated that growth after the medium exchange was more pronounced for WT cells, as likely more WT cells were removed during the medium exchange, as these cells were more planktonic. The increased propensity for attachment of the BF culture was thus highly consistent both in heterotrophic and autotrophic conditions.

Interestingly, the adapted and the WT *S. ovata* had comparable growth yields (i.e. increase of cell numbers or cell dry weight per mole of acetate produced) both under heterotrophic and autotrophic growth conditions (**Fig. 5**), meaning they have similar amounts of ATP available for growth. Biofilm formation likely comes at the expense of some ATP for the production and excretion of EPS, while acetogens already live on the thermodynamic limit of life (2). The lack of a significant difference in growth yields is possibly due to the mixed growth states of both the BF and WT cells, as none grew exclusively as a biofilm or planktonic (**Fig 4D**). In addition, the ATP required for biofilm formation can likely be saved from not expressing flagella or other traits of planktonic cells. Biofilm and planktonic cells indeed have the completely different physiological states, leading to strong differences in gene expression (13, 34, 35). *S. ovata* thus likely reallocates the small amount of energy available from acetogenesis depending on its planktonic or attached growth. Future transcriptomic and proteomic analyses could help to further map this reallocation of energy.

Overall, our results show that our ALE strategy was successful in increasing cell attachment by *S. ovata*. In general, ALE is a widely used tool for altering phenotypic traits with a poorly understood genetic basis. ALE has previously been used to successfully optimize other traits of *S. ovata* (21, 22), while it has also been used to increase biofilm formation properties of aerobic microbes (23, 24). Our study demonstrates that ALE is also a valuable method to increase attachment of anaerobic and chemolithoautotrophic microbes, such as *S. ovata*.

### Increased attachment did not lead increased performance in simple trickle bed reactors

Biofilm-based reactors systems, including trickle-bed and membrane biofilm reactors, have shown promise for H_2_:CO_2_ conversion by gas fermentation (8, 9), but so far, the importance of biofilms in these systems in not well understood. This work therefore aimed to investigate the importance of biofilm formation in biofilm-based reactor systems, by comparing the performance of the adapted *S. ovata* with that of the WT in simple trickle bed reactors. In our study, we did not find a significant difference between the performance of the two cultures in the trickle bed reactors (**Fig. 4A**), as highly similar acetate production rates and titers were obtained for both cultures, which were even slightly higher for WT *S. ovata* than for BF *S. ovata* (**Fig. 4A**). Nevertheless, an advantage for the adapted *S. ovata* was expected, as repeated medium replacements were performed to remove planktonic cells and hereby favour attachment and biofilm formation. After two of such medium replacements, higher gas consumption rates were recorded for the BF culture than for WT *S. ovata* (**Fig. 5A**), which do suggest an advantage for attached cells in the trickle bed reactors, at least for a certain period of time after the medium replacement. This advantage for attached cells clearly did not last throughout the experiment, as over the course of the experiment the proportion of planktonic cells strongly increased for both the BF and WT culture (**Fig. 4D**).

Most likely, the experimental setup used in this study was insufficient to stimulate biofilm growth. Our trickle bed reactors were simply bottles filled with carriers, which were turned around once a day to spread the medium. We did observe that carriers made the bottles function as trickle bed reactors, as the gas consumption of WT *S. ovata* was strongly increased in the presence of carriers (**Fig. 1A**). This observation was based on batch gas supply, but acetate titers with carriers were not higher than without carriers using continuous gas supply (**Fig 1B**). Under batch gas supply, the gradually decreasing gas pressures likely led to fairly low acetate production rates, as H_2_ consumption rates linearly decrease with decreasing H_2_ partial pressures (36). Those low rates likely entailed that daily trickling was sufficient to avoid to accumulation of too high acetate concentrations in the biofilms. Nevertheless, to compare the performance of the BF an WT *S. ovata*, we opted for continuous gas supply and thus a fixed H_2_ partial pressure, to avoid the kinetic effects of changing H_2_ partial pressure. Unfortunately, this continuous gas supply likely caused the daily trickling to be insufficient, as the high acetate production rates likely resulted in high local acetate concentrations in biofilms, causing product and pH inhibitions. Continuous gas supply thus likely stimulated planktonic growth, rather than biofilm formation. A proper investigation of the role of biofilms in biofilm-based reactors thus requires the use of experimental setups more resembling actual reactor operation conditions, for instance using continuously trickling and the continuous addition of new medium and removal of planktonic cells, as in more realistic, but also more complex, trickle bed reactors (9).

### Mutation in *galU* gene likely contributed to the altered biofilm formation

As the evolutionary engineering led to phenotypic changes of *S. ovata* (**Fig. 2-5**), genome comparisons were performed to investigate whether the ALE had resulted in genotypic changes. The most interesting mutation was a point mutation in the *galU* gene (change in the encoded amino acid from a leucine to a histidine (L145H)), encoding for the enzyme UTP-glucose-1-phosphate uridylyltransferase or also called UDP-glucose pyrophosphorylase (UGPase) (**Table S2**). Moreover, structural comparison of the WT and the mutated UGPase enzyme showed that this mutation possibly altered the substrate binding and the conformations of amino acid residues within the active-site pocket, leading to enhanced activity (**Fig 8**). UGPases catalyse the synthesis of UDP-glucose, which has a role in the synthesis of extracellular polysaccharides (25, 37, 38). Extracellular polysaccharides are often an important part of EPS, forming the binding matrix of microbial biofilms (14). Consequently, the identified mutation possibly resulted in the synthesis of more or more sticky EPS by the altered enzyme. Previous studies have already linked the *galU* gene to changes in biofilm formation capabilities in other bacteria, such as *Salmonella* (39), *Vibrio* (25), and *Haemophilus* (38). These studies deactivated the *galU* gene through genetically engineering and found impaired extracellular ploysaccarhirde production and in biofilm formation when the *galU* gene was deleted. Further investigations are required to verify whether the *galU* gene is also essential for biofilm formation by *S. ovata* and whether the specific point mutation found in the present study is responsible for the improved attachment of the adapted BF culture. So far, however, genetic engineering techniques to alter *S. ovata* are still in their infancy (40), but being better developed for some other model acetogens (41, 42). Once genetic engineering methods get well established for *S. ovata*, thorough investigations of the molecular mechanisms of its attachment and biofilm formation will become feasible. Moreover, comparing the performance of *WT S. ovata* with a strain deficient in biofilm formation (e.g. *GalU* deletion mutant), will form an interesting strategy to further investigate the role of biofilm formation in biofilm-based gas fermentation reactors.

## Conclusion

In this study, adaptive laboratory evolution (ALE) was applied with a selection pressure for cell attachment to enhance biofilm formation in the acetogenic bacterium *Sporomusa ovata* strain 2663. The resulting evolved culture, *S. ovata* 2663-BF, showed consistently increased cell attachment compared to the wild-type strain under both heterotrophic and autotrophic growth conditions. This biofilm-improved *S. ovata* did not have a better performance than the WT in simple trickle bed reactors, likely because the operational conditions (infrequently trickling) favored planktonic growth over cell attachment. Only after some medium replacements to remove planktonic cells, the evolved culture did have higher gas consumption rates. This evolved *S. ovata* thus offers interesting opportunities to further study the importance of biofilm formation for gas fermentation systems. The benefit of biofilm growth is likely depending on both the reactor design and operating conditions. Given *S. ovata* ‘s high performance of microbial electrosynthesis, the biofilm-improved strain is also of particular interest to unravel the role of cathodic biofilms in bioelectrochemical applications.

Genomic analysis of *S. ovata* 2663-BF revealed a notable point mutation in the *galU* gene, which encodes for the enzyme UDP-glucose pyrophosphorylase, an enzyme related to the synthesis of extracellular polysaccharides. Structural enzyme analyses suggested that this mutation likely altered the enzyme’s activity, potentially contributing to the increased cell attachment phenotype. Further research is needed to confirm that the mutation was responsible for the increased cell attachment, as well as to elucidate the broader molecular mechanisms underlying biofilm formation in *S. ovata*. Finally, the improved cell attachment did not negatively affect overall growth yields, indicating that energy for extracellular polymeric substance (EPS) production may have been reallocated from traits associated with planktonic growth.

## MATERIALS AND METHODS

### Strain and culture conditions

*Sporomusa ovata* DSM-2663 wild type (WT) was purchased from DSMZ, while *Sporomusa ovata* 2663 BF was obtained through ALE, as described in this study. Both cultures were regularly cultured at 30 °C under a N_2_:CO_2_ (80:20) atmosphere in an anoxic medium containing per L of medium 0.5 g NH_4_Cl, 0.5 g MgSO_4_ x 7 H_2_O, 0.25 g CaCl_2_ x 2 H_2_O, 2.25 g NaCl, 2 mL FeSO_4_ x 7 H_2_O solution (0.1% in 0.1 N H_2_SO_4_), 0.35 g K_2_HPO_4_, 0.23 g KH_2_PO_4_, 4 g NaHCO_3_, 0.3 g L-cysteine-HCl x H_2_O, 1 mL trace element solution (34), 1 mL selenite-tungstate solution (34) and 10 mL Wolin’s vitamin solution (DSMZ 141). Heterotrophic growth was supported by the addition of 6 g·L^-1^ betaine x H_2_O and 2 g·L^-1^ yeast extract. No betaine was added to autotrophic growth medium, which had only 0.05 g·L^-1^ yeast extract and the addition of 50 mM 3-(N-morpholino)propane sulfonic acid (pH 7). Frozen stocks (10% DMSO) were stored at −80 °C.

### Inoculation and replicates

For all experiments, the *S. ovata* cultures were revived from frozen stocks in heterotrophic growth medium and transferred once in the same medium. Cells in mid-exponential phase were used as inoculum to obtain a starting OD of 0.05. Only for trickle bed reactors under continuous gas supply, the inoculum was first washed twice with autotrophic growth medium to remove acetate and organic substrates. In addition, this inoculum was adjusted to obtain an initial cell density of about 5·10^6^ cells/mL in each reactor, as was measured using IFC. All experiments were performed at least in triplicate, while abiotic controls were included in all experiments.

### Trickle bed reactors

Simple trickle bed reactors consisted of 100 mL gas-tight bottles (with a butyl rubber stopper) filled with HDPE carriers (25 carriers with batch gas supply, 36 carriers with continuous gas supply) (high-density polyethylene plastic, HEL-X, HXF12KLL, diameter and length 12 mm, surface area 10 cm^2^, Christian Stöhr GmbH & Co., Germany). The carriers were incompletely covered with autotrophic growth medium (15 mL with batch gas supply, 10 mL with continuous gas supply). The bottles were incubated at 30°C and were regularly turned upside down and along their axis (approximately once per day, static incubation the rest of the time) to replenish nutrients and create a thin layer of adsorbed water on the carriers, simulating a trickle bed reactor.

The trickle bed reactors were operated in two different ways: either batch or continuous gas supply. Reactors with batch gas supply had a headspace consisting of N_2_:CO_2_ (80:20, 1 bar), which was over-pressurized with H_2_:CO_2_ (80:20, 0.3 bar) gas at the beginning of the experiment, as well as after the overpressure had reached zero due to microbial consumption. Gas consumption was monitored by measuring the headspace overpressure using a manometer (Bourdon).

In contrast, in trickle bed reactors with continuous gas supply, a H_2_:CO_2_ (67:33) gas mixture was continuously supplied to maintain a constant overpressure of 0.5 bar, with a maximum flow rate of 50 mL·min^-1^ as regulated by a mass flow controller (Brooks SLA5800). The liquid phase of these reactors was regularly sampled using a syringe to measure the acetate concentration, optical density (OD), and planktonic cell numbers.

### Media replacement

In the trickle bed reactors with continuous gas supply, medium was replaced to stimulate attachment and biofilm formation. To replace the medium, the bottles were gently positioned upside down to accumulate the liquid above the rubber stopper. A syringe (flushed with N_2_:CO_2_) was used to withdraw 10 mL liquid and subsequently reinject 10 mL fresh medium.

### Gas consumption rate measurements

After specific medium replacements, gas consumption rates in trickle bed reactors were measured. The continuous gas supply in these bottles was first turned off, after which the decrease in overpressure was monitored with an online manometer (Go Direct, Vernier USA). Gas consumption rates were calculated as the average decrease in pressure over time during the second hour, as the first hour was considered as equilibration time.

### Adaptive laboratory evolution

ALE was started by transferring two carriers from trickle bed reactors (with batch gas supply), initially inoculated with the original *S. ovata* 2663, to new bottles with fresh medium and fresh carriers. This transfer was performed using sterile tweezers inside an anaerobic chamber (Jacomex, France) with a N_2_:CO_2_ atmosphere (80-82% N_2_ and 18-20% CO_2_). After the transfer, a headspace pressure of N_2_:CO_2_ (80:20, 1.1 bar, to prevent underpressure) and H_2_:CO_2_ (80:20, 0.3 bar) (total pressure of 1.4 bar) was established in the new bottle. The new bottles were incubated at 30 °C and were turned upside down several times per week, while the gas consumption was monitored by measuring the decrease in headspace pressure. When the pressure approached 1 bar, H_2_:CO_2_ (80:20) gas was reinjected to reestablish a total pressure of 1.3 bar. After a second or third injection of H_2_:CO_2_ gas, the procedure of transferring carriers to a new bottle was repeated. In total, eight serial transfers were performed over a period of six months. At the later transfers, a liquid sample from the bottle (after shaking) was also taken to propagate the obtained culture in heterotrophic growth medium and prepare frozen stocks. The resulting adapted culture with increased biofilm formation is called *Sporomusa ovata* 2663-BF or BF culture in short, while the ancestral strain (*Sporomusa ovata* 2663) is called the wild type (WT) culture.

### Biofilm quantification

To assess biofilm formation characteristics, *S. ovata* (both WT and adapted BF culture) was grown in well plates (6-well plates, untreated polystyrene) with 5 mL heterotrophic growth medium per well. These well plates were placed in a plastic bag to restrict evaporation and incubated inside an anaerobic chamber (Jacomex, France) with a N_2_:CO_2_ atmosphere (80-82% N_2_ and 18-20% CO_2_) at 30 °C. Two to four days after inoculation, the well plates were taken out of the anaerobic chamber to assess biofilm formation using a crystal violet assay (13). First, the planktonic part of each well was removed and its OD_600_ was measured. Next, the biofilms remaining in the wells were stained with 5 mL 0.2% crystal violet solution for 30 minutes. After staining, the crystal violet solution was removed, and the stained biofilms were washed twice with 5 mL PBS buffer (pH 7). Finally, the stain was extracted from the biofilm with 5 mL methanol for 30 minutes, after which the absorbance of the coloured methanol solution was measured at 570 nm as a quantitative measure for biofilm formation.

### Cell number quantification

Planktonic cells numbers, as well as cells numbers attached to well plates or carriers, were quantified using impedance flow cytometry (IFC). To count the cells attached to the well plates, the planktonic phase was removed, and the remaining biofilm was carefully washed twice with PBS buffer, before the biofilm was suspended by vigorous pipetting in 5 mL fresh medium. The resulting cell suspension was agitated by vortex for 30-60 seconds in 15 mL centrifuge tubes to break up biofilms. Measured cell densities were used to calculate the number of attached cells per cm^2^ of the bottom of the wells (9.6 cm^2^).

To quantify the cell numbers attached to the carriers of the trickle bed reactors, the carriers were first washed two times inside of the reactor with autotrophic growth medium, after which all liquid was removed and the reactor was opened inside the anaerobic chamber. Seven carriers were moved carefully and with sterile tweezers to a Falcon tube to which 5 mL autotrophic growth medium was added. This tube was vortexed at high speed for at least 60 seconds, to resuspend all the cells attached to the carriers. The measured cell densities were used to calculate the average number of cells per carrier. This procedure and measurement was performed in triplicate for each reactor.

In case cell densities were not measured directly using IFC, they were estimated from the OD using the previously determined correlation factor for *S. ovata* 2663 of 2.17 · 10^8^ cells/mL per OD_600_ unit (36).

### Growth yield measurement

Growth yields of the BF and WT *S. ovata* were measured both in heterotrophic (betaine) and autotrophic (H_2_:CO_2_) growth conditions. The cultures were first grown in quintuplicate bottles with 40 mL medium and 10 carriers. For autotrophic growth, the headspace was flushed with H_2_:CO_2_ (80:20) gas and pressurized to 1.5 bar total pressure (batch gas supply). These bottles were regularly repressurized with the same gas to avoid underpressures. Once sufficient growth was observed, the bottles were vigorously shaken to detach adhered cells. A volume of 1 mL was sampled from each bottle to measure OD_600_ and cell densities using IFC. The entire liquid phase of each bottle was then filtered through a pre-dried, pre-weighed 0.25 μm polyethersulfone filter (Frisenette, DK) using a vacuum filtration system. The filter with retained biomass was dried at 55°C for 24 h, cooled down in a desiccator, and weighed on an analytical balance (± 0.0001 g precision) (VWR, LD17 4XN). Dry weight was calculated as the difference between the filter’s weight before and after filtration. In addition, the filtrate was analyzed for volatile fatty acids (VFAs). Growth yields were calculated by dividing either the dry weight or the cell numbers by the amount of acetate produced.

### Analytical methods

Impedance flow cytometry was performed using a Bactobox benchtop device (SBT Instruments A/S, Denmark) (43), placed under normal atmospheric conditions. A small volume of the homogenized cell suspensions (100-550 µL) was diluted in 10-11 mL dilute PBS buffer with a conductivity between 1900 - 2100 µS/cm and vortexed for 30-60 seconds. All reported cell numbers reflect the total particles presumed as the number of total cells in the sample. Optical densities (OD_600_) were measured at 600 nm using a spectrophotometer (Spectronic 200, Thermo Scientific).

VFAs were analyzed by GC-Flame ionization detection (FID) (system 7890, column: HP-INNOWAX 19091N-133, 30 m, 0.25 µm, 0.25 ID, Agilient Technologies, USA) with helium as the carrier gas and H_2_ and air as FID detector gases. The detector temperature was 250 °C. Pivalic acid (175 mg L^-1^ in 0.3 M oxalic acid) was used as internal standard. The detection limit for acetate was 0.3 mM. Ethanol was measure using high-performance liquid chromatography (HPLC).

### Mutation screening

The WT and BF culture were revived from frozen stocks and genomic DNA was extracted using the DNeasy Blood & Tissue Kit (Qiagen) following the manufacturer’s protocol for Gram-positive bacteria. Genomic DNA was prepared for sequencing using the Nextera XT kit (Illumina) and sequenced on an Illumina MiSeq sequencer with the MiSeq reagent kit v3. Poor quality sequences (mean <20 with a sliding window of four bases) were removed using Trimmomatic 0.39 (44). Reads were mapped to the reference sequence of *Sporomusa ovata* DSM-2662 (RefSeq accession GCF_000423685.1) and mutations were identified using Snippy 4.4. (45). Mutations found in Snippy were confirmed by assembling sequencing reads with SPAdes 3.15.5 (46) and annotation using Prokka 1.14.5 (47), checking predicted protein-coding sequences for the mutations detected by Snippy. Mutations identified in the adapted *S. ovata* 2663-BF relative to the reference sequence of *S. ovata* DSM-2662, that were not already present in the WT *S. ovata* DSM-2663, were manually inspected to check for positions relative to genes and adjacency to homopolymer regions to determine whether they were true mutations from the ALE experiment. Raw sequencing data have been deposited in the NCBI sequence read archive under BioProject accession number PRJNA1081306.

### Computational structure analysis of UGPase

The structures of the UGPase enzyme encoded by WT and the identified L145H mutant were predicted by the AlphaFold 3 server as a homotetramer (alphafoldserver.com) and subsequently filled with the product UDP-α-D-glucose and magnesium ion by the AlphaFill server (alphafill.eu) (48, 49). The final 3D structures were compared and visualized by Chimera 1.17.3 (50), while enzyme tunnel analysis was performed by Caver Web (v2.0) (51).

## Author contributions

LVG, LMD and JP designed the experiments. LVG performed the adaptation procedure and the biofilm characterization. LVG and LMD performed the trickle bed experiment with continuous gas supply. LMD quantified the growth yields and did the EPS extraction. LVG and LMD analyzed the experimental data and constructed the figures. IPGM performed the mutation analysis. NKJ and BEE performed the structural analysis of *galU*. LVG and JP wrote the manuscript. All other authors edited the manuscript. JP and KK acquired funding and JP, KK and MWVK performed supervision. All authors contributed to the article and approved the submitted version.

## Acknowledgements

This work was supported by the Novo Nordisk Foundation, as Biotechnology-based synthesis & production project grant (NNF19OC0057633). LMD was supported by a Green Transition Project I grant (MES-MODEL, 1127-00048B) from the Independent Research Fund Denmark (DFF). IPGM was supported by the Danish National Research Foundation (DNRF136). We thank Britta Poulsen, Miguel Oliviera, Simon Fruergaard and Nanna Jensen for their laboratory assistance. We also thank Heidi Skov Johansen, Thomas Dyekjær, Cecilie Nielsen, Gitte Hastrup and Linn Estel Avalon Sommer for their technical support. A thanks also to Christian Stöhr GmbH & Co. for donating the carriers applied in this study.

